# CENP-C-targeted PLK-1 regulates kinetochore function in *C. elegans* embryos

**DOI:** 10.1101/2024.04.26.591339

**Authors:** Laura Bel Borja, Samuel J.P. Taylor, Flavie Soubigou, Federico Pelisch

**Affiliations:** Molecular, Cellular and Developmental Biology, School of Life Sciences, University of Dundee. Dundee, DD1 5EH, United Kingdom

**Author notes:** Corresponding author: Federico Pelisch. ^1^ Wellcome Centre for Cell Biology & Institute of Cell Biology, School of Biological Sciences, The University of Edinburgh, Edinburgh, EH9 3BF, UK.

## Abstract

Polo-like kinase 1 (PLK1) is present in centrosomes, nuclear envelope, and kinetochores and plays a significant role in meiosis and mitosis. PLK-1 depletion or inhibition has severe consequences for spindle assembly, spindle assembly checkpoint (SAC) activation, chromosome segregation, and cytokinesis. BUB1 targets PLK1 to the outer kinetochore and, in mammals, the inner kinetochore PLK1 targeting is mediated by the constitutive centromere associated network (CCAN). BUB1-targeted PLK1 plays a key role in SAC activation and a SAC-independent role through targeting CDC-20. In contrast, whether there is a specific, non-redundant role for inner kinetochore targeted PLK1 is unknown. Here, we used the *C. elegans* embryo to study the role of inner kinetochore PLK1. We found that CENP-C, the sole CCAN component in *C. elegans* and other species, targets PLK1 to the inner kinetochore during prometaphase and metaphase. Disruption of the CENP-C/PLK1 interaction leads to an imbalance in kinetochore components and a defect in chromosome congression, without affecting CDC-20 recruitment. These findings indicate that PLK1 kinetochore recruitment by CENP-C has at least partially distinct functions than outer kinetochore PLK1, providing a platform for better understanding the different roles played by PLK1 during mitosis.

## RESULTS AND DISCUSSION

Polo-like kinases (PLKs) are a family of Ser/Thr kinases that play essential roles during the cell cycle^1^. PLKs can localise in the centrosome, nuclear envelope, and kinetochore.

Targeting to the different locations is dependent on PLK’s C-terminal polo binding domain (PBD) binding to short, phosphorylated motifs S-<u>S/T</u>-X, where the central Ser/Thr is the phosphorylated residue^2,3^. In many cases, X is a Pro residue, and this STP motif constitutes a CDK-dependent PLK binding motif.

Within the kinetochore, PLK1 plays important roles in regulating the spindle assembly checkpoint (SAC), kinetochore-microtubule attachments, and … (refs). While a plethora of PLK-binding proteins have been described to date, PLK1 kinetochore targeting is largely dependent on BUB1 in the outer kinetochore and the constitutive centromere associated network (CCAN) in the inner kinetochore^4^. In contrast to the mechanistic understanding of kinetochore targeting, little is known about the roles and substrates in the different kinetochore subpopulations. Recruited by BUB1, outer kinetochore PLK1 regulates CDC20 phosphorylation and regulates the spindle assembly checkpoint^5^ and mitosis timing^6^. In contrast, little is known about the role of inner-kinetochore PLK1. While CENP-U is the CCAN component targeting PLK1 to the inner kinetochore in mammalian cells, we have recently described that during *C. elegans* oocyte meiosis, the sole CCAN component CENP-C^HCP-4^ can target PLK-1 to different chromosome domains than BUB-1. Interestingly, the different PLK-1 populations appear to play somewhat different roles. *C. elegans* provides an excellent model system to study the mitotic division and we therefore decided to address the outstanding question on the role of inner kinetochore targeted PLK-1 using the first embryonic division.

### Inner kinetochore recruitment of PLK-1 by CENP-C

We have recently identified HCP-4, the *C. elegans* orthologue of CENP-C, as a PLK1 (hereafter ‘PLK-1’) receptor^5^. We decided to assess whether CENP-C-targeted PLK-1 plays a role during mitosis and, if so, whether this differs from that of BUB-1-targeted PLK-1^6^. Key stages of the first mitotic division of the *C. elegans* embryo are highlighted in Figure 1A for reference. We used endogenously sfGFP-tagged PLK-1^7^ to study its kinetochore recruitment and found a clear reduction in PLK-1 levels after CENP-C depletion (‘*cenp-c(RNAi)*’) at all mitosis stages (Figure 1B). Since CENP-C is required for kinetochore assembly and consequent BUB-1 kinetochore recruitment, this observation could be the trivial consequence of depleting BUB-1 from the kinetochore, which would in turn decrease PLK-1 levels^6^. We therefore took advantage of a CENP-C point mutant that disrupts its polo docking motif (‘CENP-C PD^mut^’) and cannot target PLK-1 (Figure 1C)^5^. We found that kinetochore PLK-1 levels were significantly reduced in the CENP-C PD^mut^ (Figure 1D, blue arrows). This reduction became apparent around 80 seconds after nuclear envelope breakdown (‘NEBD’, Figure 1D,E) and showed the maximal difference around metaphase, where PLK-1 kinetochore levels in CENP-C PD^mut^ were down to ∼65% (Figure 1E). This reduction was also observed in immunofluorescence of fixed samples using a specific PLK-1 antibody (Figure S1A). Importantly, abolishing PLK-1 binding to CENP-C did not affect PLK-1 at centrosomes (Figure D, yellow arrows) and did not significantly change BUB-1 levels (Figure S1B). These results suggest that CENP-C represents a parallel pathway for PLK-1 recruitment during mitosis, much like CENP-U in mammalian cells^4,8,9^.

**Figure 1.**
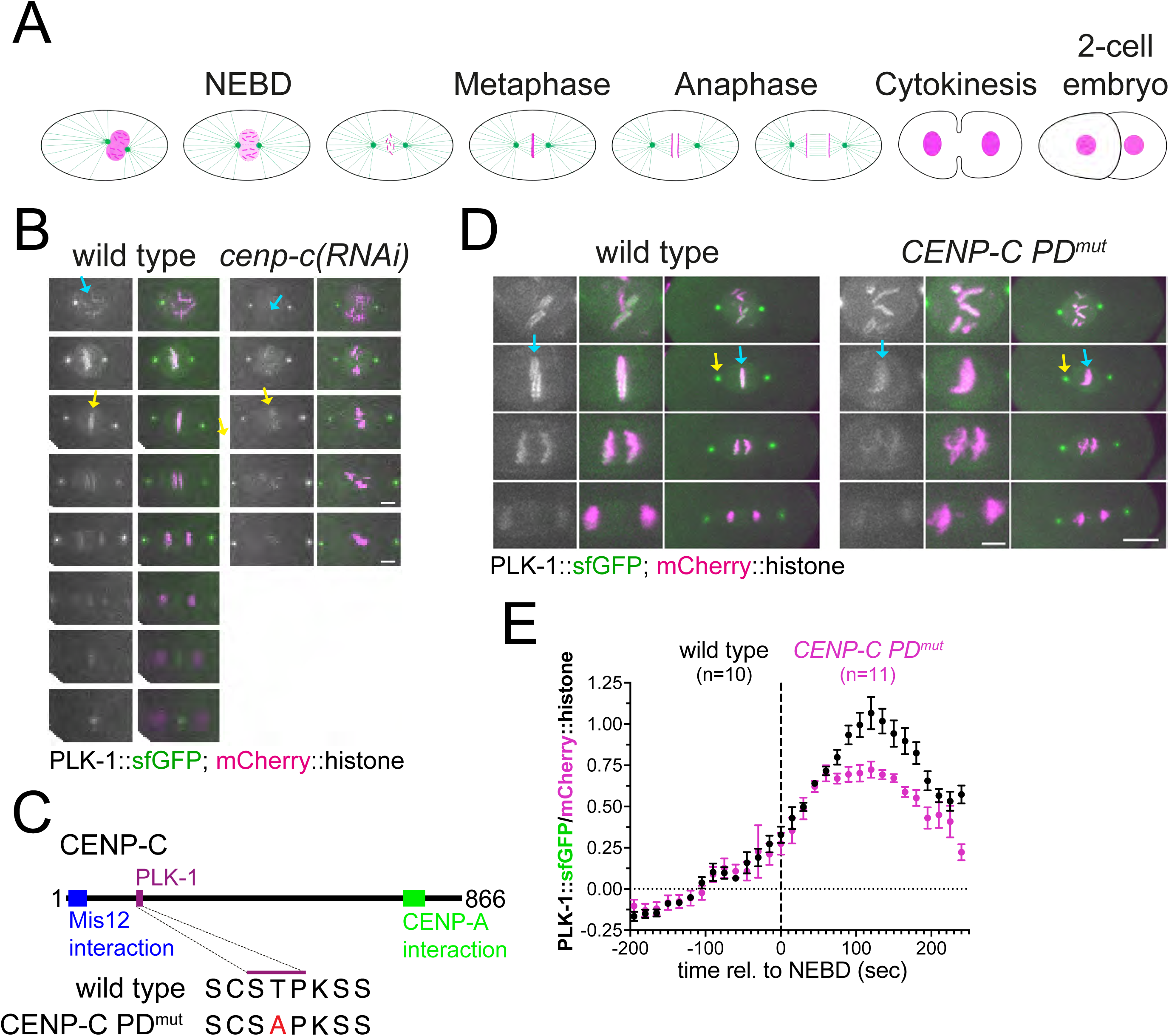
CENP-C is required for PLK-1 kinetochore localisation. **(A)** Schematic of the different stages of the first mitotic division in *C. elegans* embryos. **(B)** Fluorescently labelled PLK-1 was followed throughout mitosis in wild type and CENP-C depleted embryos (‘*cenp-c(RNAi)’*). Scale bar, 5 µm. Prometaphase and metaphase chromosomes are highlighted by cyan and yeloow arows, respectively. **(C)** Schematic indicating the PD motif characterised before^5^ along with its mutant sequence (‘CENP-C PD^mut^’), and the putative Mis12 and CENP-A interaction regions. **(D)** Fluorescently labelled PLK-1 was followed throughout mitosis in wild type and polo docking mutant CENP-C (‘*CENP-C PD^mut^’*). Scale bars, 3 µm in zoomed images and 10 µm in the whole embryo images. Yellow arrows point to the centrosome and cyan arrows to metaphase chromosomes. **(E)** Chromosomal PLK-1 levels were quantified and normalised to the chromosome signal. Values shown represent the media and the error bars denote the s.e.m. See also Figure S1.

### PLK-1 targeting by CENP-C is not involved in CDC-20 kinetochore recruitment

BUB-1-targeted PLK-1 is necessary for CDC-20 kinetochore localisation^6^. Since BUB-1 kinetochore localisation is largely unaffected in the CENP-C PD^mut^, we assessed whether inner kinetochore-targeted PLK-1 plays a different function than outer kinetochore-targeted PLK-1. We followed CDC-20 kinetochore localisation throughout mitosis and found that mutation of the CENP-C polo docking motif has no impact on CDC-20 kinetochore localisation (Figure 2A,B). To further compare the impact of disrupting CENP-C:PLK-1 binding to BUB-1:PLK-1 binding, we assessed mitotic timing, as measured by the time from nuclear envelope breakdown to anaphase onset. Whereas BUB-1 PD^mut^ displayed a longer mitosis duration as previously described before^6^, CENP-C PD^mut^ embryos displayed a slightly reduced mitotic duration (Figure 2C). These results indicate that inner kinetochore targeting of PLK-1 by CENP-C regulates different mitotic events than BUB-1-targeted PLK-1.

**Figure 2.**
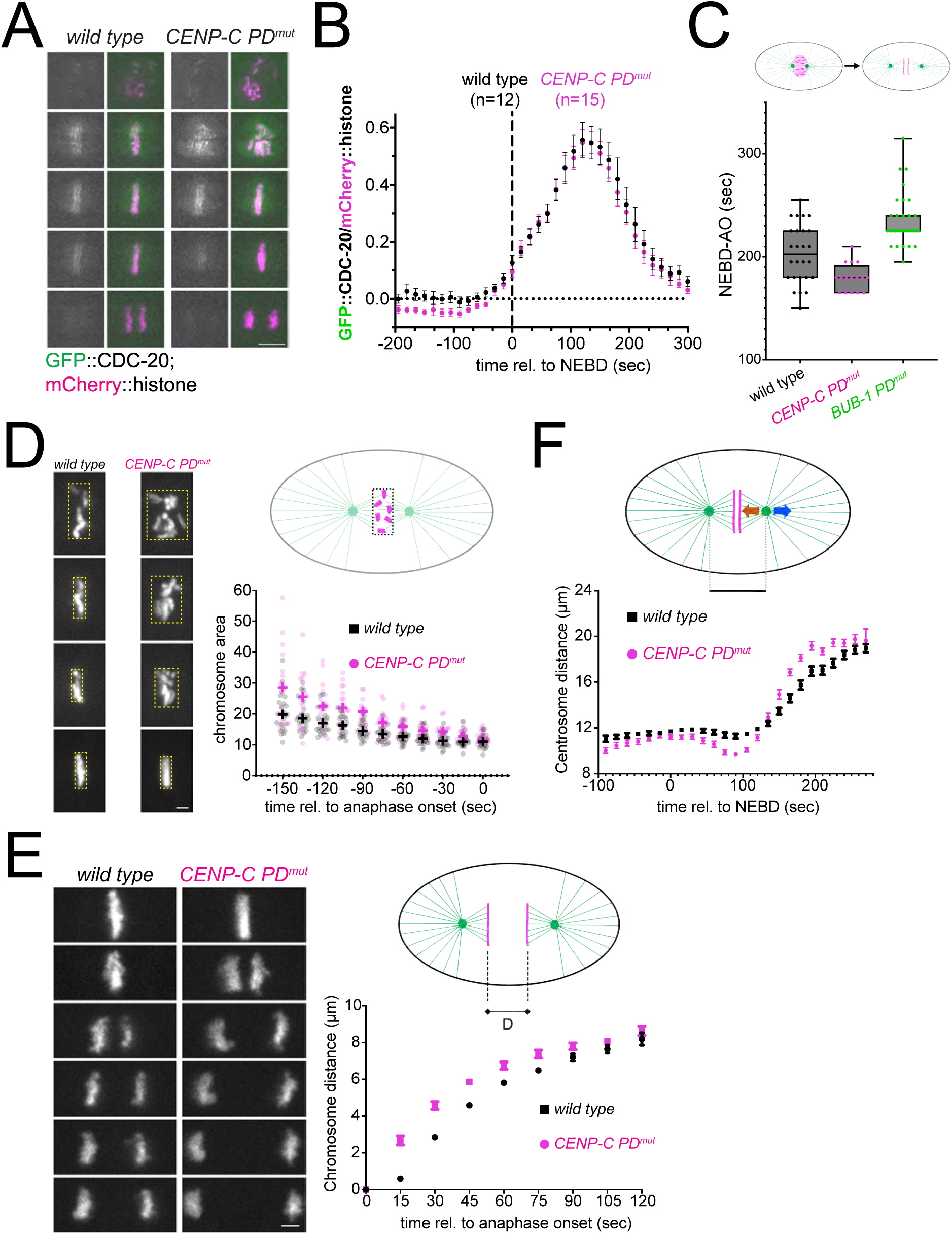
CENP-C:PLK-1 binding is not necessary for CDC-20 kinetochore recruitment. **(A)** Fluorescently labelled CDC-20 was followed throughout mitosis in wild type and polo docking mutant CENP-C (‘*CENP-C PD^mut^’*). Scale bar, 5 µm. **(B)** Chromosomal CDC-20 levels were quantified and normalised to the chromosome signal. Values shown represent the media and the error bars denote the s.e.m. **(C)** Time elapsed between NEBD and anaphase onset was measured. Circles denote the individual measurements, horizontal lines show the median, and the error bars represent the range. **(D)** Chromosome congression was assessed by bounding box analysis and the graph shows the chromosome (bounding box) area over time. Circles represent the individual measurements and the plus signs are the median for each time point. Scale bar, 2 µm. **(E)** Centrosome-to-centrosome distance was measured and the median ± s.e.m. are represented. Scale bar, 2 µm.

### PLK-1 targeting by CENP-C regulates chromosome congression and kinetochore-microtubule interactions

What mitotic events are regulated by CENP-C-targeted PLK-1? To answer this question, we chromosome congression, chromosome segregation, and kinetochore-microtubule interactions. To assess chromosome congression, we used a ‘bounding box’ method whereby we measured the area of the minimal box that surrounds all chromosomes throughout mitosis (Figure 2D). This analysis revealed that disrupting PLK-1 targeting by CENP-C leads to significant chromosome congression defects (Figure 2D). Immediately prior to anaphase onset, chromosomes do manage to congress, and the difference between wild type and CENP-C PD^mut^ becomes minimal. Our next focus was chromosome segregation during anaphase, and we measured the distance between segregating chromosomes as anaphase progresses (Figure 2E). Abolishing PLK-1 targeted BUB-1 has no appreciable effect on chromosome segregation^6^. In contrast, inhibiting PLK-1 binding to CENP-C lead to faster chromosome separation (Figure 2E). Interestingly, increased chromosome separation in the *C. elegans* first mitotic division is characteristic of kinetochore defects^10^. Prompted by this result, we decided to assess kinetochore-microtubule interactions. We measured the distance between centrosomes during mitosis, whose net movement results from the balance between astral and kinetochore microtubules (Figure 2F, blue and brown arrows on the embryo picture). Hence, the kinetics of spindle pole separation is used to infer the strength of kinetochore-microtubule attachments^11–13^. Interestingly, we noticed two significant differences between wild type and CENP-C PD^mut^ embryos. Spindle poles were significantly closer between 45 and 120 seconds after NEBD (prior to sister chromatid separation) in the CENP-C PD^mut^ (Figure 2F). This suggests that forces exerted by kinetochore microtubules are pulling stronger than the astral microtubules during this time window. The second difference was apparent during anaphase, when sister chromatids are segregating, and centrosomes begin to separate. The rate of centrosome separation was significantly higher in the CENP-C PD^mut^ (Figure 2F). As opposed to the pre-anaphase imbalance in forces, this would suggest that forces exerted by astral microtubules are not balanced by kinetochore microtubules. Altogether, the phenotypes observed in CENP-C PD^mut^ embryos suggest that CENP-C-bound PLK-1 is implicated in regulating kinetochore function.

### Inner kinetochore targeted PLK-1 regulates outer kinetochore assembly

To gain more insight into how kinetochore function is affected in CENP-C PD^mut^ embryos, we followed the dynamics of two outer kinetochore complexes: MIS12, which interacts directly with CENP-C in the inner kinetochore, and NDC80, which is present further away from centromeric chromatin and is the main point of microtubule interaction (Figure S2A). We imaged endogenously tagged KNL-3^DSN1^ during mitosis and found that loading onto chromatin started at a similar time at NEBD in both wild type and CENP-C PD^mut^ (Figure 3A,B). Immediately after NEBD, the rate of KNL-3^DSN1^ loading onto kinetochores was faster in the CENP-C PD^mut^ (Figure 3A,B), reaching also a higher maximum level compared to wild type embryos (30% higher at 165 sec post-NEBD). KNL-3^DSN1^ dissociated from chromatin during anaphase with similar dynamics in both wild type and CENP-C PD^mut^ (Figure 3A,B). CENP-C-targeted PLK-1 also regulates the MIS12 complex during female meiosis: CENP-C PD^mut^ oocytes have higher levels of kinetochore-associated KNL-3^DSN1^ (Figure 3C,D), demonstrating that kinetochore protein loading during both meiosis and mitosis is controlled by CENP-C-targeted PLK-1. Similar results were obtained in when the NDC80 complex was analysed during mitosis and meiosis (Figure S2B,C).

**Figure 3.**
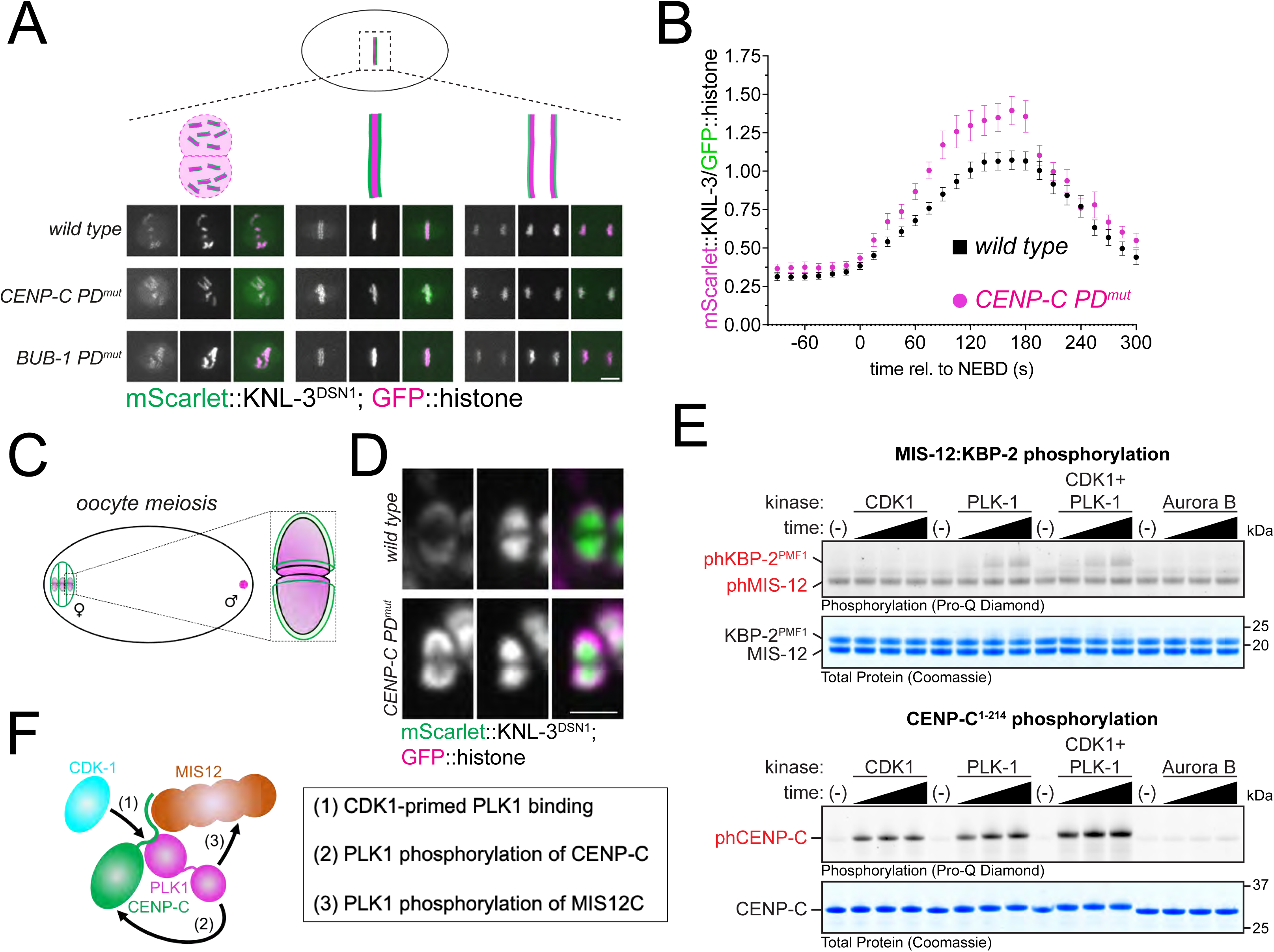
KNL-3^DSN1^ kinetochore localisation in the CENP-C PD^mut^. **(A)** mScarlet-tagged endogenous KNL-3 was followed throughout mitosis in wild type and polo docking mutant CENP-C (‘*CENP-C PD^mut^’*). Scale bar, 5 µm. **(B)** Chromosomal KNL-3 levels were quantified and normalised to the chromosome signal. Values shown represent the media and the error bars denote the s.e.m. **(C)** Schematic of an oocyte with a zoomed schematic image of a bivalent. Chromosomes are shown in magenta and kinetochores in green. **(D)** mScarlet-tagged endogenous KNL-3 was imaged during oocyte meiosis and a single image corresponding to prometaphase I is shown for wild type and polo docking mutant CENP-C (‘*CENP-C PD^mut^’*). Scale bar, 2 µm. **(E)** Kinase assays were performed with the indicated protein kinase and CENP-C or the KBP-2:MIS-12 dimer. Time points analysed were 15, 30, and 45 minutes. **(F)** Summary of the putative sequence of events leading to KBP-2 phosphorylation by PLK-1. See also Figure S2.

These results suggest that inner kinetochore PLK-1 limits the amount of outer kinetochore proteins loaded onto chromatin between NEBD and anaphase onset and could explain the greater force exerted by kinetochores in the CENP-C PD^mut^ in Figure 3C.

In mammals and Drosophila, interaction with CENP-C is mainly mediated by the MIS12:PMF1 dimer^14,15^. We purified an N-terminal fragment of *C. elegans* CENP-C^5^ and the *C. elegans* KBP-2^PMF1^:MIS-12 dimer (Figure S3) and analysed their phosphorylation status after incubation with CDK1 and PLK-1 (alone or in combination, Figure 3E). We found that whereas CDK1 phosphorylates CENP-C, as we described before, and not the KBP-2^PMF1^:MIS-12 dimer; PLK-1 phosphorylates both CENP-C and the KBP-2^PMF1^:MIS-12 dimer (Figure 3E). Interestingly, While CDK1 and PLK-1 together further enhanced CENP-C phosphorylation, no such increase was observed for the KBP-2^PMF1^:MIS-12 dimer compared to PLK-1 on its own (Figure 3E). We also tested Aurora B since it phosphorylates KNL-3^DSN1^ within the MIS12 complex^16^ but it had no impact on CENP-C or KBP-2^PMF1^:MIS-12 phosphorylation levels (Figure 3E). A plausible model stemming from these results is that CDK1 phosphorylates CENP-C in the inner kinetochore driving PLK-1 recruitment. PLK-1 would then phosphorylate CENP-C and KBP-2^PMF1^ (among other putative substrates) to regulate kinetochore function (Figure 3F). However, further work will be required to characterise the landscape of PLK-1 substrates.

### Disruption of inner and outer kinetochore PLK-1 leads to severe defects in mitotic chromosome segregation

Having established that inner kinetochore targeted PLK-1 plays different roles than outer kinetochore PLK-1 during mitosis, we decided to assess the consequences of inhibiting PLK-1 recruitment by both BUB-1 and CENP-C. Chromosome alignment and segregation were severely affected by the combined mutation of PLK-1 docking sites (Figure 4A). We decided to focus or further analysis on chromosome segregation (Figure 4B,C) and found that 100% of CENP-C PD^mut^/BUB-1 PD^mut^ embryos displayed abnormal segregation, which included mild (52%) and severe (48%) defects (Figure 4C). Interestingly, while the combined CENP-C and BUB-1 PD mutants displayed lower PLK-1 levels than BUB-1 PD^mut^ (Figure 4D, P=0.025 Mann Whitney Test), the difference was not significant compared to CENP-C PD^mut^ alone (Figure 4X, P=0.25 Mann Whitney Test). These results show that the severity of the phenotypes do not correlate with the quantity of PLK-1 and therefore suggest that a qualitative difference (i.e. substrates). It should be noted that some PLK-1 is still detectable in kinetochores, which could be due to 1) the little amount of endogenous BUB-1 remaining in the BUB-1^T527A^ mutant condition (see methods) and/or 2) the existence of another PLK-1 centromere/kinetochore receptor. In summary, joint disruption of the main inner and outer kinetochore platforms for PLK-1 recruitment leads to severe mitotic defects, reminiscent of the joint depletion of CENP-U and BUB1 in mammalian cells^17^.

**Figure 4.**
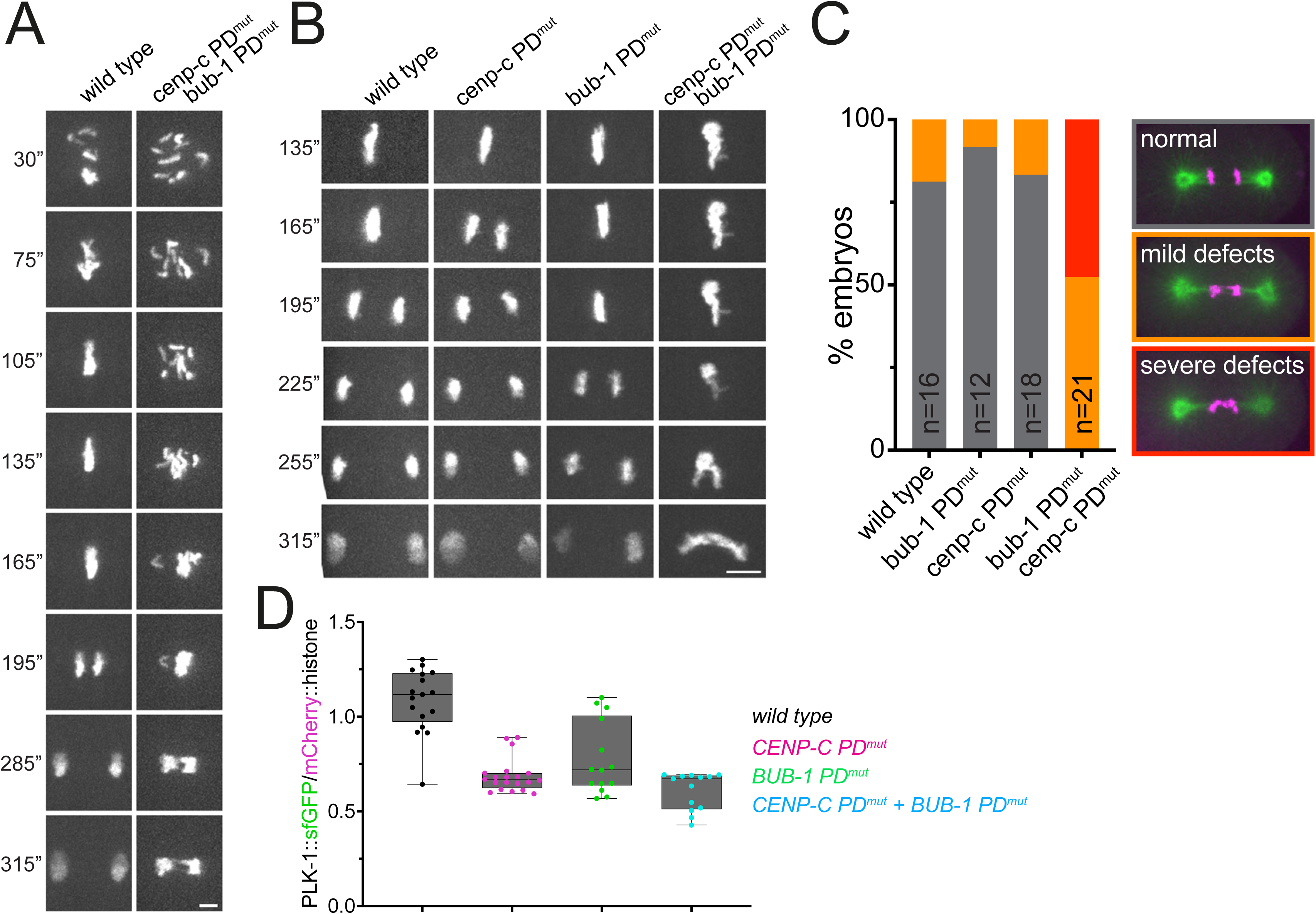
KNL-3^DSN1^ kinetochore localisation in the CENP-C PD^mut^. **(A)** GFP-tagged histone was followed throughout mitosis in wild type and the double polo docking mutant *CENP-C PD^mut^/BUB-1 PD^mut^*. Times are relative to NEBD (0”). Scale bar, 3 µm. **(B)** GFP-tagged histone was followed throughout mitosis in wild type and the double polo docking mutant *CENP-C PD^mut^/BUB-1 PD^mut^*. Times are relative to NEBD (0”). Scale bar, 5 µm. **(C)** Chromosome segregation defects were arbitrarily divided into mils and severe (See Methods) and the percentage of embryos displaying any phenotype for each condition was quantified. n denotes the number of embryos analysed for each condition. **(D)** Chromosomal PLK-1 levels were quantified and normalised to the chromosome (mCherry::histone) signal. Values shown represent the median and the error bars denote the range. Individual points (single embryos) are shown. Quantitation corresponds to 135 seconds post-NEBD. See also Figure S3.

We have provided evidence supporting a specific role for inner kinetochore-bound PLK-1 during mitotic chromosome segregation. Interestingly, CENP-C-bound PLK-1 appears to play a more important role than CENP-U in mammals since abrogation of CENP-C:PLK-1 binding leads to kinetochore-microtubule attachment defects as well as chromosome congression defects. In mammalian cells, these effects are seen mainly when the CENP-U:PLK-1 binding is abrogated in cells not expressing BUB1^17^. Our experiments using point mutants instead of depletion to abolish PLK1 binding to BUB-1 and CENP-C show that there is still PLK-1 remaining on kinetochores, suggesting the existence of other pathway(s) targeting PLK-1 to kinetochores. However, this remaining kinetochore PLK-1 population is not sufficient to sustain proper chromosome segregation. We also provide additional insight by describing that MIS12 and NDC80 complexes increase at kinetochores in the absence of inner kinetochore-bound PLK-1. It will be critical to identify inner and outer kinetochore PLK-1 substrates to understand the mechanisms underlying PLK-1’s different functions. Additionally, we are also focusing in identifying additional PLK-1 receptors.

## ACKNOWLEDGEMENTS

We thank Arshad Desai for sharing strains and antibodies. This work was supported by a Career Development Award from the Medical Research Council (grant MR/R008574/1) and an ISSF grant funded by the Wellcome Trust (105606/Z/14/Z). ST was funded by a Medical Research Council Doctoral Training Programme. We acknowledge the help and support of the Dundee Imaging Facility. Some nematode strains were provided by the CGC, which is funded by NIH Office of Research Infrastructure Programs (P40 OD010440).

## AUTHOR CONTRIBUTION

L.B-B. conducted some imaging experiments. F.S. conducted immunofluorescence experiments and commented on the manuscript. S.T. expressed all recombinant proteins used in the manuscript and conducted the in vitro kinase assays. F.P. designed the experiments, performed some experiments, and performed the data analysis.

### SUPPLEMENTAL INFORMATION

Figures S1-S3 and Table S1.

### SUPPLEMENTAL FIGURE LEGENDS

**Figure S1.**
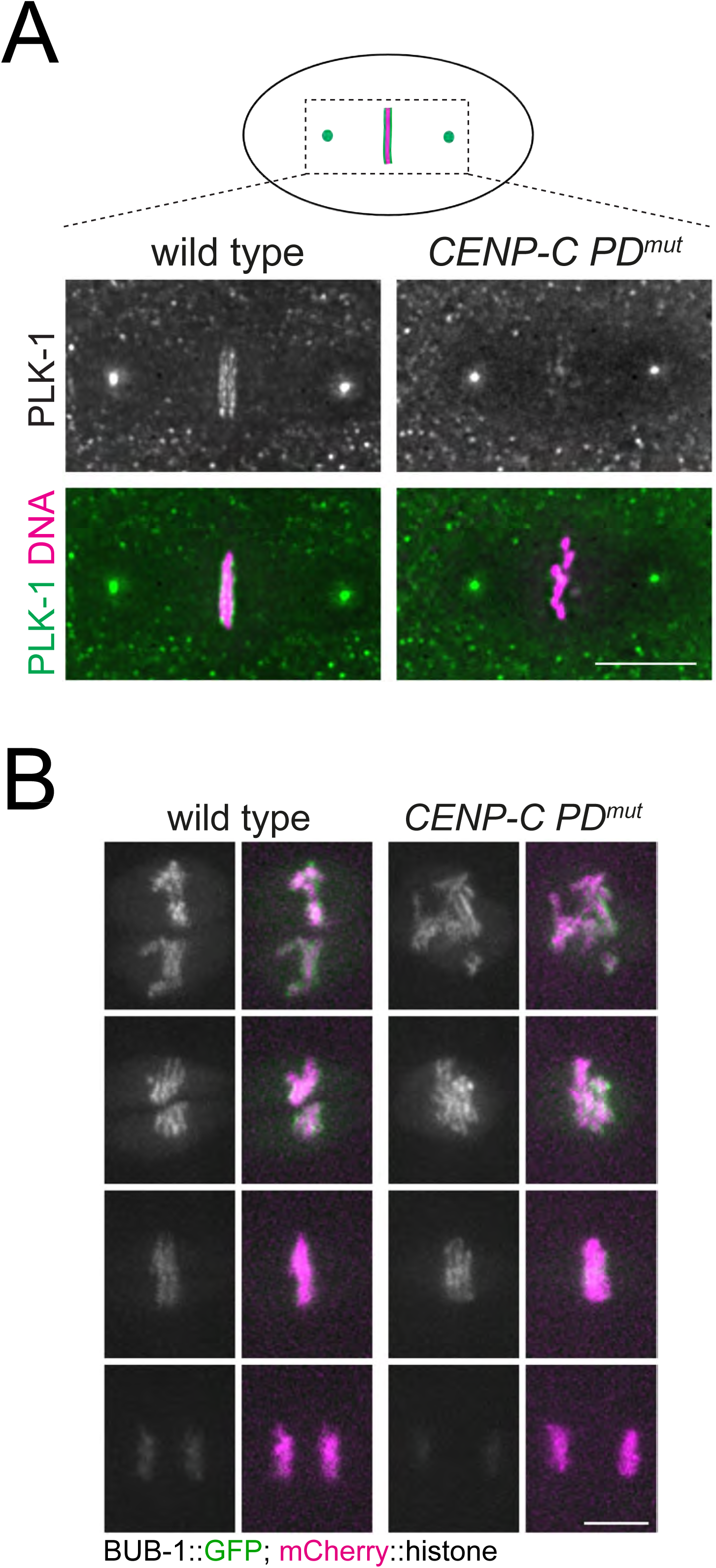
CENP-C is required for PLK-1 kinetochore localisation. **(A)** mmunofluorescene analysis using a specific anti-PLK-1 antibody was carried out in wild type and CENP-C PD^mut^ embryos at metaphase. Scale bar, 5 µm. **(B)** Fluorescently labelled BUB-1 was followed throughout mitosis in wild type and CENP-C PD^mut^ embryos. Scale bar, 5 µm.

**Figure S2.**
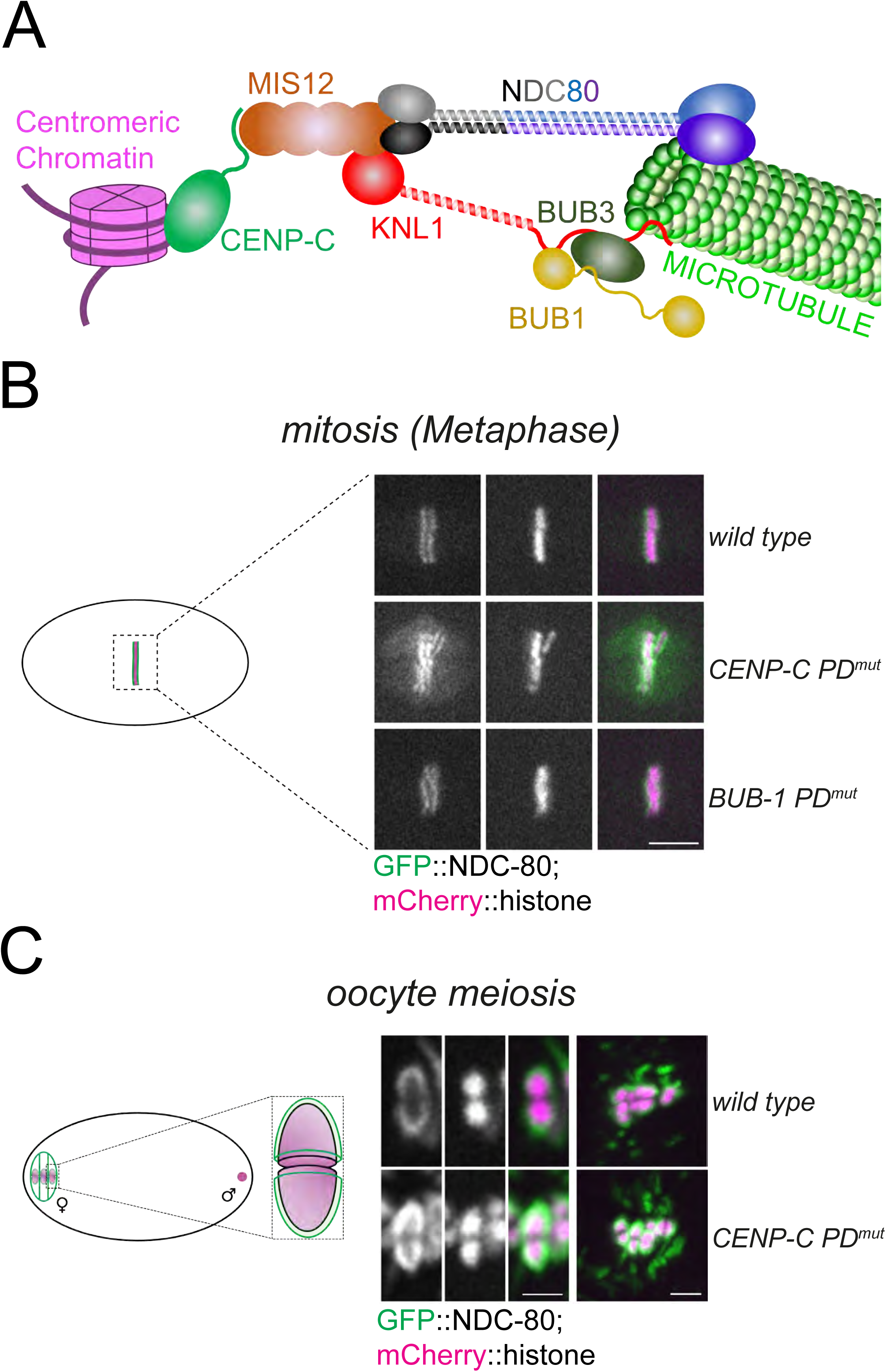
NDC-80 kinetochore localisation in the CENP-C PD^mut^. **(A)** Schematic depicting how CENP-C connects the centromerric chromatin with the outer kinetochore, consisting of the KMN network (KNL-1/Mis12 complex/NDC80 complex) and the BUB complex (BUB-1/BUB-3). NDC-80 complex provides the major point of contact with microtubules. **(B)** GFP-tagged endogenous NDC-80 was followed throughout mitosis in wild type, CENP-C PD^mut^, and BUB-1 PD^mut^. Scale bar, 5 µm. **(C)** GFP-tagged endogenous NDC-80 was imaged during oocyte meiosis and a single image corresponding to prometaphase I is shown for wild type and polo docking mutant CENP-C (‘*CENP-C PD^mut^’*). Scale bars, 2 µm in the zoomed panel (left) and 3 µm in right panel.

**Figure S3.**
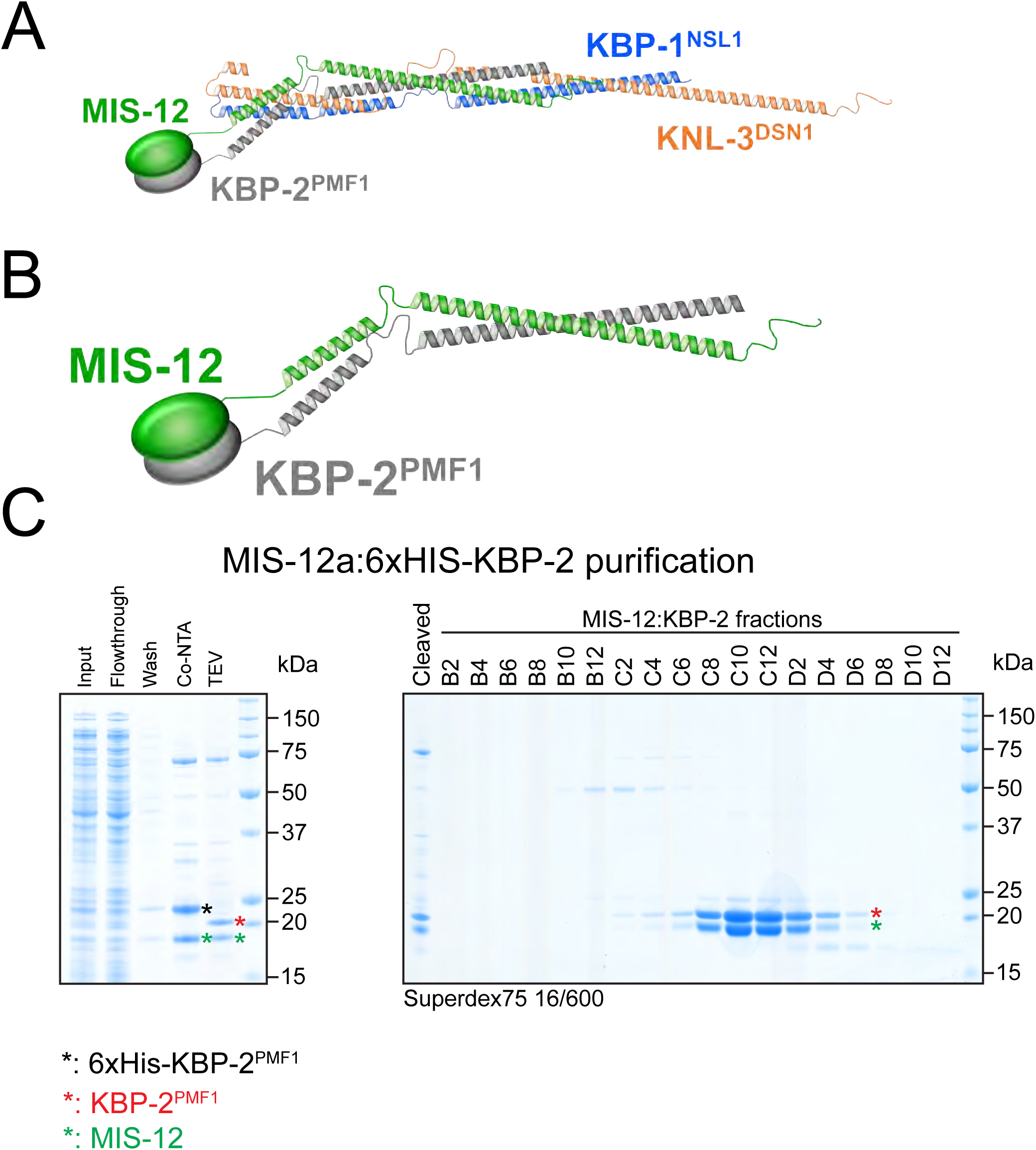
Purification of the MIS-12:KBP-2^PMF1^ dimer. **(A)** Schematic of the full MIS-12 complex structure based on an AlphaFold2 prediction. The disordered region of KNL-3^DSN1^ (aa 1-159) is not shown. **(B)** Enlarged image of the MIS-12:KBP-2^PMF1^ complex from (A). **(C)** Biochemical purification of the MIS-12:KBP-2^PMF1^ complex from bacteria.

## METHODS

### *C. elegans* strains & RNAi

Strains used in this study were maintained at 20 degrees unless indicated otherwise. For a complete list of strains, please refer to Table S1.

For RNAi-mediated depletions, the targeting sequence for *bub-1* was 2353-2935 and for *hcp-4*, 967-2128, both from the first ATG codon. For double depletion, both sequences were cloned in the same vector. All sequences were inserted into L4440 using the NEBuilder HiFi DNA Assembly Master Mix (New England Biolabs) and transformed into DH5α bacteria.

The purified plasmids were then transformed into HT115(DE3) bacteria^18^. RNAi clones were picked and grown overnight at 37°C in LB with 100 μg/ml ampicillin. Saturated cultures were diluted 1:100 and allowed to grow until reaching an OD600 of 0.8–1. Isopropyl-β-D-thiogalactopyranoside (IPTG) was added to a final concentration of 1 mM, and cultures were incubated for 1 h at 37°C. Bacteria were then seeded onto NGM plates made with agarose and 1 mM IPTG and allowed to dry. L4 worms were then plated on RNAi plates and maintained at 20°C.

### Live imaging of embryos

Embryos were dissected and mounted in 5 µl of L-15 blastomere culture medium (0.5 mg/mL Inulin; 25 mM HEPES, pH 7.5 in 60% Leibowitz L-15 medium and 20% heat-Inactivated FBS) on 24×40 mm #1.5 coverslips. After dissection and isolation of early embryos, a circle of vaseline was laid around the sample, and a custom-made 24×40 mm plastic holder (with a centred window) was placed on top. The sample was imaged immediately using 488 nm and 561 nm laser lines. Live imaging was done using a CFI Plan Apochromat Lambda 60X/NA 1.4 oil objective mounted on a microscope (Nikon Eclipse Ti) equiped with a Prime 95B 22mm camera (Photometrics), a spinning-disk head (CSU-X1; Yokogawa Electric Corporation). Acquisition parameters were controlled with NIS software (Nikon). For all live imaging experiments, partial projections are presented. All files were stored, classified, and managed using OMERO^19^. Figures were prepared using OMERO.figure and assembled using Adobe Illustrator. Representative movies shown in Supplementary material were assembled using Fiji/ImageJ^20^ custom-made macros.

In Figure 3A, the strains used were FGP739, FGP740, and FGP741, all in the prescence of *bub-1(RNAi)* to deplete endogenous BUB-1. In Figure 3D, the strains used were FGP722 and FGP728. In Figure S2B, mitotic analysis was performed using strains FGP803, FGP804, and FGP805, all in the prescence of *bub-1(RNAi)* to deplete endogenous BUB-1. In Figure S2C (meiotic analysis) FGP372 and FGP729 were used.

### Kinase Assays

Reactions were carried out using 55 nM CDK1/CyclinB, 75 nM PLK-1, or 200 nM Aurora B in 50 mM Tris-HCl, pH 7.5; 1 mM ATP; 10 mM MgCl_2_; 0.5 M TCEP; 0.1 mM EDTA.

Substrates were used at ∼16 µM (CENP-C) or ∼10 µM (KBP-2/MIS-12 dimer). Reactions were conducted at 30°C for the indicated time points. Samples were taken and added to an equal volume of 2X LDS buffer (Thermo) and incubated at 70°C for 15 min or 95°C for 5 min before loading onto SDS-PAGE gels.

### Immunofluorescence

Worms were placed on 4 µl of M9 worm buffer in a poly-D-lysine (Sigma, P1024)-coated slide and a 24×24-cm coverslip was gently laid on top. Worms were dissected to release the embryos, slides were placed on a metal block on dry ice for >10 min. The coverslip was then flicked off with a scalpel blade, and the samples were fixed in methanol at 20°C for 30 min. Secondary antibodies were donkey anti–sheep, goat anti-mouse, or goat anti-rabbit conjugated to Alexa Fluor™ 488, Alexa Fluor™ 594, and Alexa Fluor™ 647 (1:1,000, Thermo Scientific). Donkey anti-mouse and donkey anti-rabbit conjugated secondary antibodies were obtained from Jackson ImmunoReserach. Embryos were mounted in ProLong Diamond antifade mountant (Thermo Scientific) with DAPI. Primary antibodies were: α-PLK-1^21^, α-HCP-4 ^13^, α-BUB-1, purified in house after immunisation of rabbits using the sequence in^12^,

### Protein Intensity Measurments

A semi-automated Fiji macro was used to measure the levels of the protein of interest (POI). Briefly, a ‘sum slices’ projection was generated for each data point and the signal in the histone channel was taken as a reference. This selection was then transferred to the POI channel and the intensity value was taken. All graphs were generated using Graphpad Prism 10.

### Chromosome Congression Analysis

The maximum intensity projection in the mCherry::histone channel was used to analyse chromosome area, using the bounding box selection tool in Fiji^20^. Graphs were prepared using Graphpad Prism 10.

### Chromosome Segregation Defects

Chromosome segregation defects were quantified in two arbitrary categories: 1) ‘mild defects’, when either lagging chromosomes or minimal anaphase bridges were detected and 2) ‘severe defects’, when either severe anaphase bridges or the chromosome masses remained as a single body during anaphase. The graph was created with Graphpad Prism 10.

---

**Table S1:** *C. elegans* strains used in this study.

|  |  |  |  |
| --- | --- | --- | --- |
| bub-1::linker::gfp; mCherry::histone | FGP202 | *bub-1(syb1134[bub-1::linker::gfp])I; ltIs37 [pAA64; pie-1/mCherry::his-58; unc-119 (+)]IV; unc-119(ed3)III* | This study |
| PLK-1::sfGFP; mCherry::histone | FGP263 | *plk-1(lt18[plk-1::sGFP]::loxP)III; ltIs37 [pAA64; pie-1/mCHERRY::his-58; unc-119 (+)]IV* | doi.org/10.1016/j.devcel.2017.09.019 |
| NDC-80::GFP; mCherry::histone | FGP372 | *ndc-80(lt54[ndc-80::GFP::tev::loxP::3xFlag])IV; ltIs37 [pAA64; pie-1/mCHERRY::his-58; unc-119 (+)]IV* | This study |
| hcp-4(T163A) | FGP669 | *hcp-4(fgp58[hcp-4(T163A)])I* | doi.org/10.7554/eLife.84057 |
| PLK-1::sGFP; mCherry::histone; hcp-4(T163A) | FGP719 | *plk-1(lt18[plk-1::sGFP]::loxP)III; ltIs37 [pAA64; pie-1/mCHERRY::his-58; unc-119 (+)]IV; hcp-4(fgp58[hcp-4(T163A)])I* | doi.org/10.7554/eLife.84057 |
| mScarlet::KNL-3; GFP::histone | FGP722 | *knl-3(dha19 [mScarlet-I<sup>3</sup>XFLAG::knl-3] V; ruIs32 [pie-1::GFP::histone + unc-119(+)], unc-119 (ed3)* | This study |
| mScarlet::KNL-3; GFP::histone; hcp-4(T163A) | FGP728 | *knl-3(dha19 [mScarlet-I<sup>3</sup>XFLAG::knl-3] V; ruIs32 [pie-1::GFP::histone + unc-119(+)], unc-119 (ed3); hcp-4(fgp58[hcp-4(T163A)])I* | This study |
| NDC-80::GFP; mCherry::histone; hcp-4(T163A) | FGP729 | *ndc-80(lt54[ndc-80::GFP::tev::loxP::3xFlag])IV; ltIs37 [pAA64; pie-1/mCHERRY::his-58; unc-119 (+)]IV; hcp-4(fgp58[hcp-4(T163A)])I* | This study |
| mScarlet::KNL-3; GFP::histone; bub-1(wt) | FGP739 | *knl-3(dha19 [mScarlet-I<sup>3</sup>XFLAG::knl-3] V; ruls32 [pie-1::GFP::histone + unc-119(+)], unc-119 (ed3); unc-119(ed3)III;ltSi268[pOD/pTK013; Ppub-1::Bub1 reencoded; cb-unc-119(+)]II* | This study |
| mScarlet::KNL-3; GFP::histone; hcp-4(T163A); bub-1(wt) | FGP740 | *knl-3(dha19 [mScarlet-I<sup>3</sup>XFLAG::knl-3] V; ruls32 [pie-1::GFP::histone + unc-119(+)], unc-119 (ed3); hcp-4(fgp58[hcp-4(T163A)])I; unc-119(ed3)III;ltSi268[pOD/pTK013; Ppub-1::Bub1 reencoded; cb-unc-119(+)]II* | This study |
| mScarlet::KNL-3; GFP::histone; bub-1(T527A) | FGP741 | *knl-3(dha19 [mScarlet-I<sup>3</sup>XFLAG::knl-3] V; ruls32 [pie-1::GFP::histone + unc-119(+)], unc-119 (ed3); unc-119(ed3)III;ltSi1346[pTK070;Ppub-1::BUB-1 T527A reencoded; cb-unc-119(+)]II* | This study |
| NDC-80::GFP; mCherry::histone; bub-1(wt) | FGP803 | *ndc-80(lt54[ndc-80::GFP::tev::loxP::3xFlag])IV; ltIs37 [pAA64; pie-1/mCHERRY::his-58; unc-119 (+)]IV; Ppub-1::Bub1 reencoded; cb-unc-119(+)]II* | This study |
| NDC-80::GFP; mCherry::histone; hcp-4(T163A); bub-1(wt) | FGP804 | *ndc-80(lt54[ndc-80::GFP::tev::loxP::3xFlag])IV; ltIs37 [pAA64; pie-1/mCHERRY::his-58; unc-119 (+)]IV; hcp-4(fgp58[hcp-4(T163A)])I; unc-119(ed3)III;ltSi268[pOD/pTK013; Ppub-1::Bub1 reencoded; cb-unc-119(+)]II* | This study |
| NDC-80::GFP; mCherry::histone; bub-1(T527A) | FGP805 | *ndc-80(lt54[ndc-80::GFP::tev::loxP::3xFlag])IV; ltIs37 [pAA64; pie-1/mCHERRY::his-58; unc-119 (+)]IV;ltSi1346[pTK070;Ppub-1::BUB-1 T527A reencoded; cb-unc-119(+)]II* | This study |
| bub-1::linker::gfp; mCherry::histone; hcp4(T163A) | FGP815 | *bub-1(syb1134[bub-1::linker::gfp])I; ltIs37 [pAA64; pie-1/mCherry::his-58; unc-119 (+)]IV; unc-119(ed3)III; hcp-4(fgp58[hcp-4(T163A)])I* | This study |
| GFP::CDC-20; mCherry::histone | OD2591 | *ltSi814[pPLG047; Pfzy-1::gfp::fzy-1::fzy-1 3'UTR; cb-unc-119(+)]I; unc-119(ed3)III?; ltIs37 [pAA64; pie-1/mCHERRY::his-58; unc-119 (+)] IV* | doi.org/10.1101/gad.302067.117 |
| GFP::CDC-20; mCherry::histone; hcp-4(T163A) | FGP829 | *ltSi814[pPLG047; Pfzy-1::gfp::fzy-1::fzy-1 3'UTR; cb-unc-119(+)]I; unc-119(ed3)III?; ltIs37 [pAA64; pie-1/mCHERRY::his-58; unc-119 (+)] IV; hcp-4(fgp58[hcp-4(T163A)])I* | This study |

